# Extrinsic mortality and senescence: a guide for the perplexed

**DOI:** 10.1101/2022.01.27.478060

**Authors:** Charlotte de Vries, Matthias Galipaud, Hanna Kokko

**Affiliations:** Department of Evolutionary Biology and Environmental Studies, University of Zurich, Zurich, Switzerland; Department of Biological and Environmental Science University of Jyväskylä, Jyväskylä, Finland; Institute for Biodiversity and Ecosystem Dynamics, University of Amsterdam, The Netherlands; Konrad Lorenz Institute of Ethology University of Veterinary Medicine Vienna, Austria; Faculty of Biological and Environmental Sciences University of Helsinki, Helsinki, Finland; Institute for Organismic and Molecular Evolution, Johannes Gutenberg-University Mainz, Mainz, Germany

**Keywords:** Senescence, Life-History Evolution, Trade-Offs, Fast-slow life histories, Density dependence

## Abstract

Do environments or species traits that lower the mortality of individuals create selection for delaying senescence? Reading the literature creates an impression that mathematically oriented biologists cannot agree on the validity of George Williams’ prediction (who claimed ‘yes’). The abundance of models and opinions may bewilder those that are new to the field. Here we provide heuristics as well as simple models that outline when the Williams prediction holds, why there is a ‘null model’ where extrinsic mortality does not change the evolution of senescence at all, and why it is also possible to expect the opposite of William’s prediction, where increased extrinsic mortality favours slower senescence. We hope to offer intuition by quantifying how much delaying the ‘placement’ of an offspring into the population reduces its expected contribution to the gene pool of the future. Our first example shows why sometimes increased extrinsic mortality has no effect (the null result), and why density dependence can change that. Thereafter, a model with ten different choices for population regulation shows that high extrinsic mortality favours fast life histories (Williams) if increasing density harms the production of juveniles or their chances to recruit into the population. If instead increasing density harms the survival of older individuals in a population, then high extrinsic mortality favours slow life histories (anti-Williams). We discuss the possibility that empirically found Williams-like patterns provide indirect evidence for population regulation operating via harming the production or fitness prospects of juveniles, as opposed to the survival of established breeders.

## Introduction

> “It is not the case that additional mortality automatically favours the evolution of senescence.”
>
> — Caswell and Shyu, 2017

> “Reports of the death of extrinsic mortality moulding senescence have been greatly exaggerated.”
>
> — Jack da Silva, 2018

> “Williams’ intuition about extrinsic mortality is irrelevant”
>
> — Moorad et al. 2020b

The above quotes lay bare a rather odd state of affairs: more than six decades after Williams (1957) presented his argument for the relationship between adult mortality rates and the evolution of senescence, mathematically trained biologists still cannot seem to agree on what patterns theory actually predicts. Williams’ seminal work argued that populations experiencing different rates of mortality (as adults) should senesce at different rates (Gaillard & Lemaître 2017). The intuitive message is that if life is bound to be short ‘anyway’ (due to, e.g., high predation risk), it makes little sense to invest in a robust body able to resist ‘wearing out’ for a long time (Medawar 1952, Williams 1957).

William’s work has since been interpreted to mean that an increase in age-independent extrinsic mortality — typically defined as the result of hazards from the environment which are constant throughout life (Koopman et al. 2015, see Moorad et al. 2020a for definitional issues) — should select for faster senescence (Da Silva 2018, Dańko et al. 2018, André and Rousset 2020). Others have argued against this idea, stating that age-independent extrinsic mortality cannot affect the evolution of senescence (Gadgil and Bossert 1970, Taylor et al. 1974, Abrams 1993, Caswell 2007, Caswell and Shyu 2017, Wensink et al. 2017, Moorad et al. 2019). Recent work, while aiming to clarify, has simultaneously led to a large number of different models and opinions, which as a whole may be confusing to those that are new to the field (André and Rousset 2020, Dańko et al. 2017,2018, and the debate started by Moorad et al. 2019 and continued in Day & Abrams 2020, da Silva 2020, Moorad et al. 2020a,b). Here our aim is to explain what happens in models of senescence when extrinsic mortality increases. Specifically, we aim to outline when the prediction made by Williams holds and when we can expect it to fail.

In the following, we call, for the sake of conciseness, the prediction that age-independent extrinsic mortality does not impact senescence ‘the null result’. The null result can be interpreted to mean that ‘Williams was wrong’, but it is useful to distinguish the null from an even stronger way for a prediction to disagree with the Williams hypothesis: it is logically possible that higher extrinsic mortality associates with *slower* (not faster) senescence (Abrams 1993). Thus, we have a range of potential results which we, for brevity, call ‘Williams’ (is right), ‘null’, and ‘anti-Williams’.

The null result is typically explained using selection gradients (e.g. Caswell 2007, Caswell & Shyu 2017). Selection gradients generally measure how much fitness changes (on average) when a trait takes a different value from the current population mean. In the current context, the relevant traits are life history traits such as survival or fecundity. The null result refers to a situation where the selection gradient associated with survival takes the same value across all possible values of extrinsic mortality. This is a surprising result for anyone whose intuition aligns with Williams. Yet it arises not only when selection gradients are computed in models that lack density dependence, but also when there is density dependence that impacts survival of all ages equally (Caswell 2007). A related but alternative derivation of the null result, and of deviations from the null-result, can be found in the appendix of Day & Abrams’ (2020) which uses growth rate optimization to quantify the effect of an increased extrinsic mortality under different kinds of density dependence. Here we hope to provide an intuitive explanation for the null result by instead focusing explicitly on the time that a newborn is placed into a population. Delaying the ‘placement’ of an offspring into the population reduces its expected contribution to the gene pool of the future — but only if a population is growing.

Our work below has two parts. The first part aims to create the simplest possible setting where selection for senescence can be stronger or weaker. We strive for simplicity by ignoring many real-life complications, such as trade-offs between survival and reproduction, as this allows us to assume that the contrasted populations only differ in extrinsic mortalities, not e.g. in fecundities. Also, in the first part we keep the life cycles very simple: reproduction can happen either once or twice. This setting already contains sufficient ingredients where the null result can remain intact or be invalid, depending on what we assume about population regulation (which stops unlimited exponential growth). Next, we move to a second set of models, where we add realism by modelling senescence with Gompertz-Makeham survival curves (Gompertz 1825, Makeham 1860). Gompertz-Makeham survival curves are commonly used in the field of demography: these functions assume that mortality has an intrinsic component that increases exponentially with age (Missov and Lenart 2013). We introduce life-history trade-offs by assuming that an organism can avoid this increase if it accepts a lower rate of reproduction. We examine the outcomes of this trade-off under a range of different types of population regulation. This exercise shows the choice of regulation can flip systems from the ‘null’ pattern to either ‘Williams’ or ‘anti-Williams’. In other words, ‘fast’ lives (in the sense of fast and slow life histories, Stearns 1989, Promislow and Harvey 1990), may evolve as a response to high extrinsic mortality, but this outcome does not happen under every form of population regulation.

Our second approach also allows linking senescence to the ‘understudied territory’ identified by Moorad et al. (2019): what happens when a population does not stabilize to zero growth but fluctuates, so that there are years (or, more generally, time steps) with increasing and others with decreasing population sizes (see also Caswell & Shyu 2017)? Fluctuations in population abundance due to continually occurring stochastic fluctuations in the vital rates are a common way to model such situations (Tuljapurkar 2013, Caswell & Shyu 2017), but populations might also fluctuate due to events that occur less often and cause large mortality in a pulsed manner, a scenario that we include. These events may impose age-dependent or stage-dependent mortality. A population may be regulated via these events if they happen more often at high density (e.g. a disease spreads), and the population may then spend much of its time growing towards high density rather than remaining near an equilibrium (in other words, transient dynamics become important). In this case predictions based on selection gradients might not apply (Capdevila et al. 2020), since their calculation requires demographic stability or small stochastic and age-independent fluctuations around a demographic equilibrium (Caswell and Shyu 2017).

In our examples below, density dependence that acts on fecundity has a different impact on senescence than density dependence that acts on survival (across all ages). While these results are fully in line with previous insights (Abrams 1993, Caswell and Shyu 2017, Wensink et al. 2017, Dańko et al. 2017,2018, and other papers cited above), we hope that our presentation will make the issues more heuristically transparent.

### An example free of tradeoffs: why does the null result arise?

Being able to fly is often quoted as an example of reduced extrinsic mortality (Austad and Fischer 1991, Holmes and Austad 1994, Healy et al. 2014). Although this is clearly not the only reason for e.g. bat lifespans exceeding those of similarly sized rodents (for complexities, e.g., hibernation, see Wilkinson and Adams 2019), we take the dichotomy ‘volant or not’ as a way to conceptualize extrinsic mortality differences in our first, trade-off-free model. We ask whether a bat, assumed to experience relatively low extrinsic mortality, will be selected more strongly to delay senescence than a similar-sized non-volant organism, such as a mouse. Note that ignoring life-history trade-offs means that we are, in this first exercise, not interested in the fact that litter sizes are smaller in bats than in rodents; we wish to consider the effect of mortality in isolation of everything else. Reproductive effort and its potential trade-offs with senescence will be considered in the second part of our paper (see also the appendix of Day & Abrams 2020 for an analytical example with trade-offs).

We further simplify the situation (away from real life, but helpful for heuristic understanding) by assuming a finite lifespan that does not permit more than one or two breeding attempts. Both the bat and the mouse have two alternative life histories that differ in their rates of senescence in a simple and dramatic fashion: a ‘fast-senescer’ can only breed once and always dies thereafter, while a ‘slow-senescer’ can breed up to two times (we also include survival up to each breeding event). Clearly, both mice and bats will benefit from adding an extra breeding attempt to their lifespan. Selection for a second breeding attempt is therefore positive for both species, and the key aim is to compare the *strength* of selection for the two species. If bats benefit much more from the extra breeding attempt than mice, then selection on bats to reduce senescence is stronger and the result is in line with Williams’ hypothesis.

Each mouse individual survives with probability *s*_M_ from birth to first reproduction, and slow-senescing mice additionally have the same probability of surviving after their first breeding to reach their second attempt. For bats, the rules are the same, but the survival probabilities equal *s*_B_. Since we assume all else is equal, we assign the same fecundity *F* to mice and to bats. *F* also does not change between the first and the second breeding attempt. Since there are already many analytical results available (e.g. Day & Abrams 2020), and our aim is to aid intuition maximally, we will make use of a single numerical example where *s*_B_=3 s_M_, i.e., bat survival is three times that of mice, and we show results assuming 20% survival in mice, 60% in bats (Table 1 gives an overview of bat and mice life-history parameters).

**Table 1:**
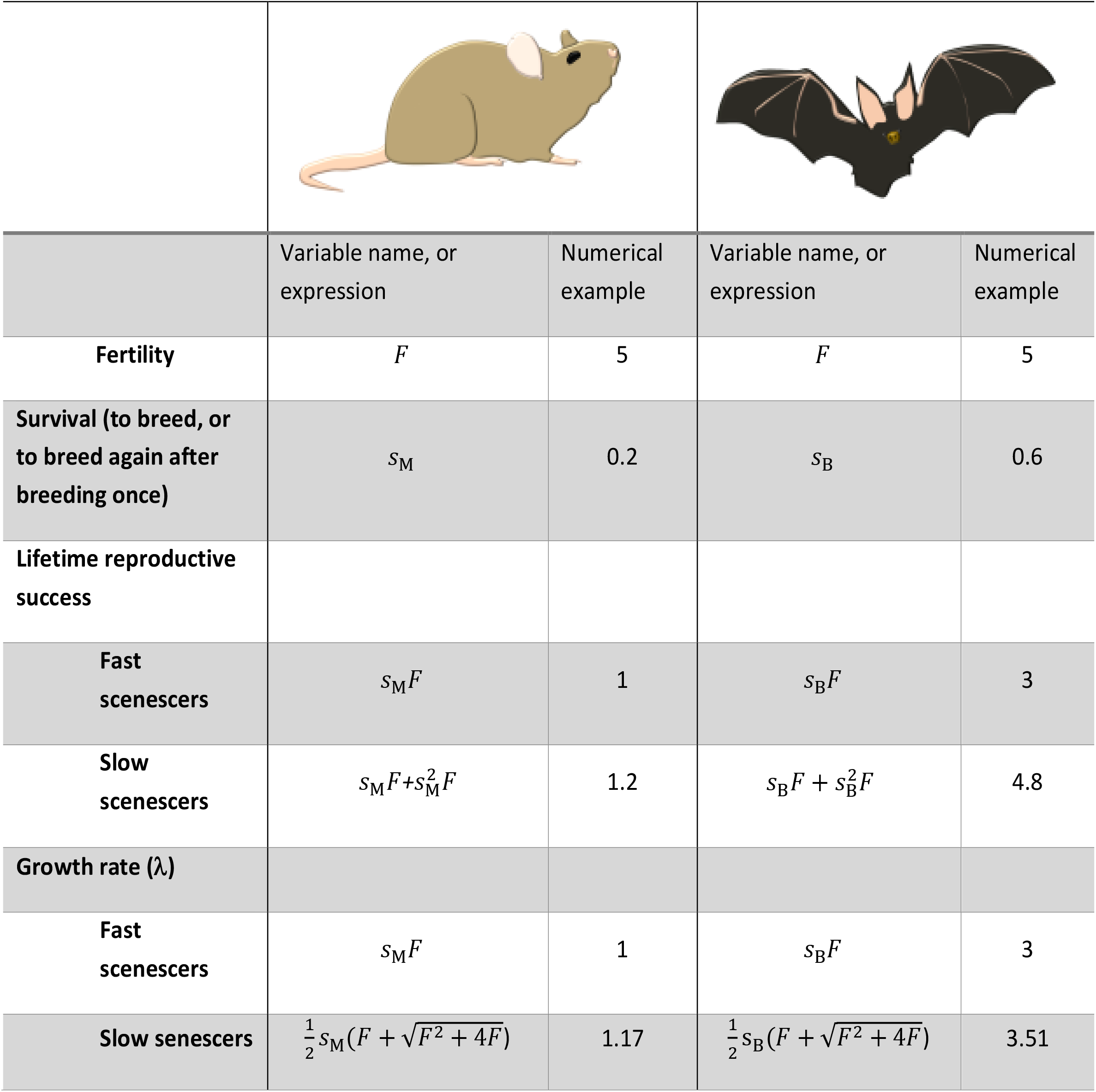
List of variables and their values used in the trade-off free example. To calculate λ, start from the Euler-Lotka equation for a species with ω age classes,

|  |  |  |  |  |
| --- | --- | --- | --- | --- |
|  | Variable name, or expression | Numerical example | Variable name, or expression | Numerical example |
| **Fertility** | $F$ | 5 | $F$ | 5 |
| **Survival (to breed, or to breed again after breeding once)** | $s_M$ | 0.2 | $s_B$ | 0.6 |
| **Lifetime reproductive success** |  |  |  |  |
| **Fast senescers** | $s_M F$ | 1 | $s_B F$ | 3 |
| **Slow senescers** | $s_M F + s_M^2 F$ | 1.2 | $s_B F + s_B^2 F$ | 4.8 |
| **Growth rate ($\lambda$)** |  |  |  |  |
| **Fast senescers** | $s_M F$ | 1 | $s_B F$ | 3 |
| **Slow senescers** | $\frac{1}{2} s_M (F + \sqrt{F^2 + 4F})$ | 1.17 | $\frac{1}{2} s_B (F + \sqrt{F^2 + 4F})$ | 3.51 |

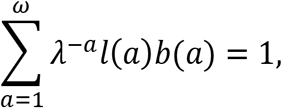

where *l*(*a*) is the proportion of individuals surviving until age *a*, and *b*(*a*) is the mean reproductive output of these survivors. For example, for the slow senescing mouse the Euler-Lotka equation becomes

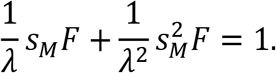

Solving for λ, gives the equation in the table.

The lifetime reproductive success (LRS) of slow-senescing bats is increased by 4.8/3 = 1.6 relative to the fast-senescing bats, i.e. an improvement of 60%. The LRS of slow-senescing mice is increased by 1.2/1 = 1.2 relative to fast-senescing mice, i.e. an improvement of 20%. It is not a coincidence that 60% and 20% are identical to the survival values we assigned to the two species since 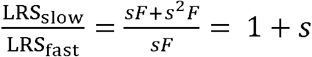, thus s is a direct measure of the expected improvement over the baseline. Since the improvement in LRS of bats was a factor three times the improvement in LRS of the mice when gaining the ability to breed twice, one might be tempted to conclude that bats are selected to reap the benefits of a long life much more strongly than mice, based on the extrinsic mortality argument (*s*_B_ > *s*_M_).

However, this conclusion is premature, and this illustrates a key argument in the debate. In the absence of density regulation, the superior survival of bats compared with mice also makes their population grow much faster than that of mice – in our example, their growth rate is precisely threefold (Table 1). This result applies for any positive value of *F*: the terms containing *F* in the calculation of the growth rate λ are identical for bats and for mice. It does, however, require that bat fecundity does not differ from mouse fecundity, which is simply a reminder that we are focusing here on the effect of extrinsic mortality alone, and leave fecundity considerations for later.

An important point to note here is that LRS is only a valid fitness measure if density dependence acts on fecundity of individuals of all ages equally (Mylius and Dieckmann, 1995). In the absence of any density dependence, populations will be growing exponentially and the population with the fastest growth rate will dominate. In general, invasion fitness is the only reliable fitness measure (see Kokko 2021 for a review about population fitness), but under some circumstances invasion fitness simplifies to a familiar life history measure such as the population growth rate, or the life-time reproductive success (see discussion in Mylius and Dieckmann, 1995). Intuitively, if two individuals both have the same LRS but one produces its offspring earlier, these (and their descendants) form a larger proportion of the future gene pool in a growing population.

To quantify precisely how important it is to reproduce early in a fast growing population, it is useful to calculate how much early produced offspring contribute to the total population at some later time *t*, and contrast this with a late produced offspring, a procedure we do both for the mouse and the bat population. Since this calculation requires both types of offspring (early and late) to exist, we focus on an initial parent that is, necessarily, a slow-senescer, and also assumed to be among the lucky ones who survive to breed twice. The offspring themselves are examples of slow-senescing life histories with the appropriate survival rates. These initial founding offspring, of which we consider 1 each (early and late produced) in both species, are placed into a population that is growing exponentially at the appropriate species-specific rate (Figure 1, with growth rates from Table 1). The differing timing of offspring placement into the population is graphically illustrated as an earlier and a later star symbol in Figure 1), and since the populations of both mice and bats are assumed to grow, the late-produced offspring form a smaller proportion of the population at the time of production than the early-produced offspring. This initial disadvantage has consequences ever after. Measured at a later time point, the proportion of the population that descends from the early-produced offspring is far larger than the proportion descending from the late-produced offspring in the (well-surviving and hence fast growing) bat population. This difference also exists in the mouse population, but it is much less extreme in this species (the widths of the two ‘stripes’ denoting lineages show only moderate differences in Figure 1a, and strong differences in Figure 1b).

**Figure 1.**
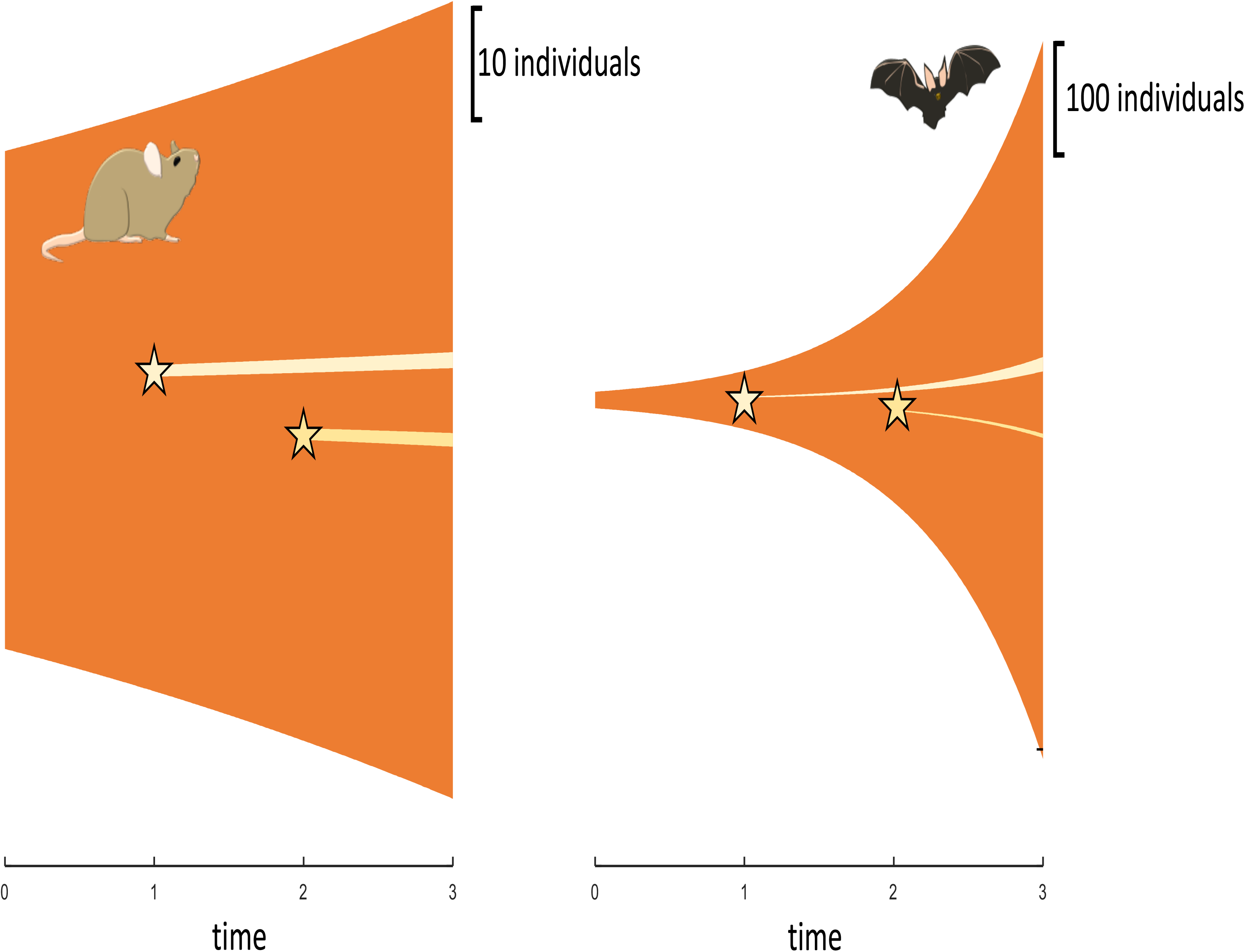
A visualization of why the bat, with its threefold growth rate, is penalized more for a delay in placing offspring into a growing population. For purely aesthetic reasons, population growth is depicted over continuous time (*r* = ln(λ)). Values of *r* correspond to λ values given in Table 1. Populations are assumed to consist of slow senescers (Table 1) and stars depict the placement of one offspring at time *t* = 1 or *t* = 2 into a growing population. Both species consist of 50 individuals at time *t* = 1 (shown at a different scale as indicated, to fit the entire growth into picture, as bats, with their higher survival, increase their numbers much faster than mice. For both species, the lineage (pale stripes) that starts with an offspring placed into the population at *t* = 2 is thinner than the lineage that had its start at *t* = 1, but this difference is much more marked if population growth is (b) fast than if it is (a) slow. In (b) both stripes appear narrow, because the vertical scale has to differ between (a) and (b) to allow the entire bat population to be depicted.

These differences can be quantified. The descendants of an early-produced offspring, *N*_B,early_, as well as a late-produced offspring, *N*_B,late_, will eventually reach a stable proportion of young and ‘old’ (namely second year) individuals, forming two lineages that both grow at a rate identical to the population growth rate (Caswell 2001). It follows that the lineage arising from the early-produced offspring, measured at some time *t* after both lineages have been initiated, is larger than the lineage arising from the population of descendants of the late-born offspring, by a factor of *λ*_B_. That is,

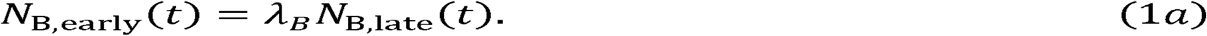

Likewise for the descendants of the early and late offspring of a focal mouse individual,

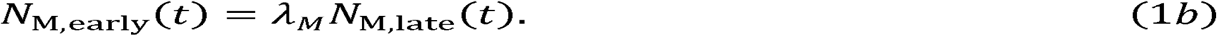

If we divide both sides by the total population (this total contains additionally all other descendants from this or other parents), we obtain the proportion of the total population at time *t* that represents descendants of early and late offspring, respectively. The proportion of the population descended from the original bat parent’s early offspring is larger than that of her late offspring by a factor *λ*_B_, and for mice, this is *λ*_M_. Since 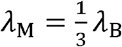 (Table 1), the early produced bats are worth three times more (relative to their later produced siblings) than early produced mice (relative to theirs). In other words, in a population that is growing at a threefold rate compared with another, the importance of reproducing early is also elevated by the exact same factor (threefold). This is analogous to investing money into a growing economy: the faster the growth, the better off are those who were able to invest early; the penalty (discounting) of late investments is visible in Figure 1 as the trumpet shaped pale stripes (descendant lineages) being more unequal in height for the bat than for the mouse.

Therefore, we have a situation where on the one hand it appears more ‘profitable’ to have the ability to breed twice if chance often permits this longevity to really materialize (in the bats), but on the other hand, this very ability allows populations to grow fast, and this means that late-produced offspring are, to borrow an economic term, discounted (much less valuable). The cancelling out reflects the fact that one could argue both ‘for’ and ‘against’ bats being the species that experiences stronger selection to survive to breed twice. The argument for those who root for the bats: clearly selection for a robust body that can breed twice only pays off if extrinsic circumstances allow this to be materialized, and extrinsic circumstances do so three times more often for bats than for mice. The equally appealing counterargument is that late-produced offspring are a particularly poor investment in bats, as the good survival of all individuals means that a late-produced young forms a much smaller proportion of the gene pool than an early-produced one. In the mice, this penalty is only a third of what it is for bats. The net truth is that both arguments are valid, but since they are valid simultaneously, the factors (3 and 1/3) cancel out and bats experience exactly as strong selection to breed twice as do mice.

In our numerical example (Table 1), the growth rate increases by 17% when shifting from being a fast-senescer to a slow-senescer. Importantly, this number applies whether one is a bat or a mouse (from table 1: for mice, 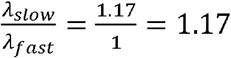 and for bats, 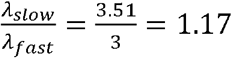). The equally strong growth rate improvement is the reason behind the statement that selection to become a slow senescer is the same in bats and mice, despite the latter being exposed to much higher mortality (lower *s*) throughout their lives. The example works with other *s*_B_ : *s*_M_ ratios too, which can be seen by dividing the expressions in the last row of Table 1 with those on the penultimate row. The values of s_B_ and s_M_ simply cancel out, and the values λ_slow_/λ_fast_ become identical for the two species.

To conclude, even though being able to delay senescence until after the 2^nd^ breeding attempt (instead of dying after the 1^st^) benefits bat LRS much more than mice if surviving to breed is more likely for bats, LRS fails as a predictor of selection because it does not take into account that late-produced offspring are also less valuable than the early-produced ones — and this decline in value is much faster for the species that, by virtue of its high survival, has faster population growth. Since we assumed that higher survival directly translates into a higher growth rate, the ‘penalty’ of placing offspring late into the population is far greater for bats than for mice. These two effects (better improvement in LRS, and the larger penalty) cancel each other out exactly. The outcome is the ‘null result’: selection for slow life-histories (against senescence) is equally strong in the bat and the mouse population.

This result can also be confirmed by comparing population growth rates of entire populations of fast-senescers versus slow-senescers. Calculating the population growth rate improvement of slow-senescing bats and mice relative to their fast-senescing competitors yields the same answer for both species: both improvements are 17% (with data from Table 1, note that 1.17 / 1 = 3.51 / 3). Since population growth rate is the correct fitness proxy for exponentially growing or declining populations (Charlesworth, 1994; Mylius and Diekmann, 1995; Caswell, 2001), not the LRS, this section has confirmed that age-independent extrinsic mortality does not affect the relative benefit of reduced senescence for species experiencing different levels of extrinsic mortality, *in the absence of density dependence*.

### Beyond the null: what cancels out under density dependence, what does not?

Above, we intentionally considered an unrealistic comparison, to be able to show what happens if survival is the only difference between two populations. Real bat populations do not show threefold growth compared with mice, and neither can sustain exponential growth forever. Intuition (to some at least) suggests that the slow-senescing bats can begin to truly reap the benefits of a long life if density dependence makes the ‘penalty’ of having to discount the value of late-produced offspring less severe. Why? If the population does not in reality expand as fast as predicted by density-independent growth rules “5 offspring per year and 60% survival for all who aren’t scheduled to die of old age yet”, then the trumpet shapes of Figure 1 do not expand as fast as they did before. Mathematically, slower growth means that the value of late-placed offspring is not devalued as strongly compared to the early-placed ones. Therefore, as a whole density dependence offers a potential for a smaller penalty for a lineage of descendants appearing late into a population. If population growth ceases altogether, the penalty vanishes as well. In other words: *if we assume that slow bats can reach old age just as often as they did in the density-independent case*, and now their late offspring are not nearly as bad investments as they were under unlimited population growth, then selection is now much freer to reward slow life histories (Figure 2 illustrates the idea graphically).

**Figure 2.**
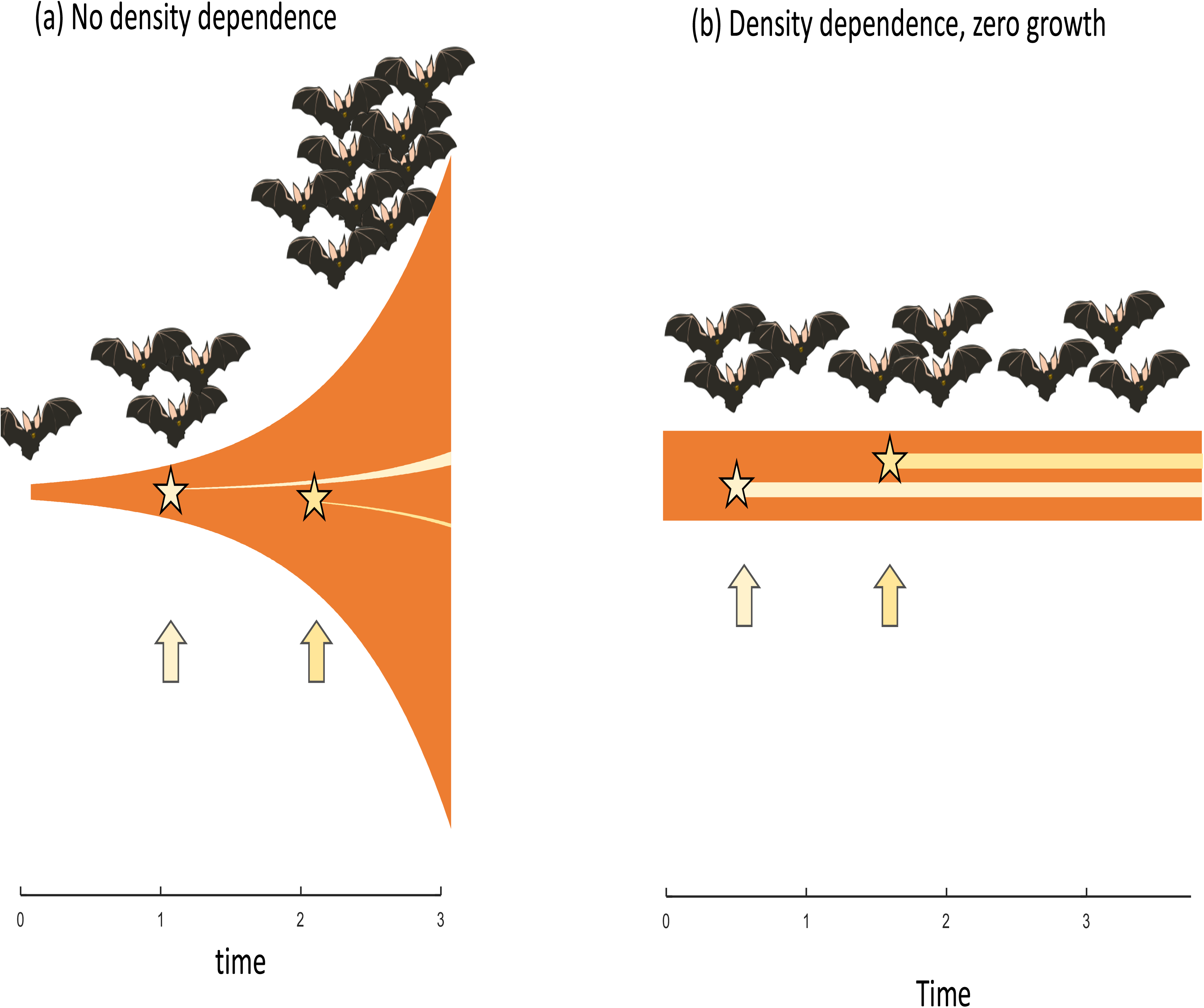
A visualization of the difference between placing offspring early or late in an exponentially growing population (on the left) or in a stable population (on the right). In a growing population, the late placed offspring will be a smaller fraction of the future population than the early placed offspring. In a stationary population, there is no such discounting of early *vs*. late placed offspring.

This intuition can be correct, but it comes with a strong caveat: the *if* clause in the previous sentence. The argument relies on the assumption that bats really can reach old age just like they did under unlimited growth. The crux of the issue is that population growth cannot be reduced ‘just like that’; that is, while keeping all vital rates unchanged. Something has to change for the growth to be lower. Perhaps fewer young are born, or perhaps some are never born because their potential mothers had already died. There are many possibilities, and this matters.

If slowed down population growth is achieved by making reproduction somewhat harder for everyone, then it is indeed possible that the chances that a slow-senescing bat reaches old age remain the same (*s*_B_^2^ = 0.36 in our example) across all densities (Abrams 1993, Day & Abrams 2020). In this situation, the slowing down and eventual stabilization of population growth can begin favouring delaying senescence in those organisms that are relatively likely to reach old age in the first place (i.e. bats as opposed to mice in our example). Slow-senescing mice, too, enjoy some of this advantage, but only 4% of them do, because high extrinsic mortality (*s*_M_^2^ = 0.04) means most (96%) do not live to enjoy their intrinsic ability to breed twice.

But, importantly, slowing down (the tendency of *r* = ln(λ) to decline towards 0) can also be achieved via other mechanisms. High densities could, for various ecological reasons, make it very hard for older bats (or mice) to survive while the fecundity of survivors is left intact. Now it is quite hard to be convinced that those who *in principle* have good prospects for reaching old age (bats, as opposed to mice, in our example) would also *in practice* achieve this benefit. If density regulation effectively prevents slow-senescing types from translating their intrinsic survival ability to actual survival (and subsequent reproduction), selection will be blind to their slow-senescing phenotypes.

This can explain why the ‘null’ result sometimes happens even when density dependence is included (e.g. Caswell and Shyu 2017). Typically, in these cases, a range of extrinsic mortality values are compared between hypothetical populations, but each population is also forced to have zero growth (*r* = 0). If the condition *r* = 0 is achieved by adjusting mortality rates at all ages equally, then, effectively, the initial elevation of extrinsic mortality (for those populations in the comparison who were supposed to have high extrinsic mortality) is removed again from the model by density-dependent adjustments of the mortality itself. Some people argue that this is a fundamental and exciting proof that helps us understand why extrinsic mortality cannot matter (Moorad et al. 2019); Moorad et al. 2020a make their preference for including the total effects of a mortality adjustment more explicit still. Others might reason that this particular exercise is somewhat pointless, as it assumes that underlying variations in extrinsic mortality will not be visible in the mortality schedules experienced by individuals at equilibrium. Phrased in the context of our example, they would never be measurable as real bats having lower mortality than real mice.

Because this example is important, we repeat the message in the context of an experiment. While our example is hypothetical, it is inspired by experimentally imposed high and low adult mortality regimes in *Drosophila* populations (Stearns et al. 2000). Imagine that an empiricist aims to cause high mortality in the regime where this is the intention, but ends up realizing that the individuals that were not removed by the experimenter respond with improved survival; this is possible since they now live in a less dense population. If the high-mortality regime first increases mortality and then, by reducing competition among the survivors, lowers it, it is possible that total mortality remains unchanged. Any measures of senescence rates remain unchanged as well. Did the researcher recover a deep insight, confirming Moorad et al’s message? Or will she instead respond by stating “my experiment didn’t work - it remained uninformative because the manipulation failed to produce an actual difference in the mortality actually experienced by the population, making the subsequent finding that senescence didn’t change a trivial one”? We leave it to the reader to form their own opinions about this matter, as we believe both viewpoints have their merits. It is of interest to note that Moorad et al. 2020a identify a difference in Day & Abrams’ (2020) thinking compared to theirs based on whether the label ‘extrinsic mortality’ is applied before or after various consequences, such as those in our hypothetical experiment, have been allowed to act on the population.

To sum up, by now, we have achieved some intuition as to why it is important to identify who precisely fails to survive, or fails to be born, when increasing densities reduce population growth. The key question is: can a slow-senescing phenotype reap the benefits of its long life across all relevant population densities, or are its survival prospects themselves affected by density? If survival of older individuals is left relatively intact and so is the value of late-placed young (due to the population no longer growing so fast), then we can expect the Williams prediction to hold. If the slow-senescer, on the other hand, itself suffers from density increases because survival of old slow-senescers is disproportionally targeted by density regulation, then we may enter the realm of the null, or even an anti-Williams region (see Abrams 1993 for examples).

### Ten case studies of slow and fast life histories

To make our thought experiment above as simple to follow as possible, we focused on an ‘all else being equal’ comparison where the two species did not differ in fecundity and there were no trade-offs: an ability to delay senescence required no lowering of reproductive effort. We next turn to examples that are considerably more realistic than the above comparison between hypothetical species that only differ in one respect (survival) and cannot ever breed more than twice. We now sacrifice analytical tractability to achieve three goals: (i) we consider a wide variety of density-dependent scenarios; (ii) we link senescence to the ideas of fast and slow life histories (which is argued to underlie e.g. the mammal-bird dichotomy in senescence rates, Jones et al. 2008, relationships between senescence and latitude across bird species, Møller 2007, all the way to within-species patterns, Cayuela et al. 2020), taking into account that a slow senescence rate may involve ‘accepting’ lower fecundity; and (iii) we see if the intuition remains robust in situations (identified as important by Moorad et al. 2019) where populations fluctuate around an equilibrium due to pulsed high mortality events instead of staying invariant or experiencing small, stochastic fluctuations around the equilibrium (Tuljapurkar 2013, Caswell & Shyu 2017).

We explore ten different kinds of density regulation, of which nine are organized in a 3×3 setup (Table 2) and an additional one (density dependence acting on recruitment probability) added for the reason that this form of population regulation is often discussed in territorial species (Newton, 1992; Sæther et al. 2002; López-Sepulcre and Kokko, 2005; Krüger et al. 2012; Grant et al. 2017). In the 3×3 scheme, we have three examples each of density dependence acting on (1) survival in an age-independent manner, (2) on adult survival (neither the number of juveniles nor adults impacts juvenile survival) or (3) on fecundity (noting also that fecundity regulation in this case is mathematically indistinguishable from newborns dying, or having trouble recruiting into the population; see Discussion). Each of these is investigated in three different ways: density dependence may be absent for a while until it acts in a pulsed (‘catastrophic’) manner via sudden decreases in the vital rates either (a) deterministically above a certain density or (b) stochastically, or (c) density dependence may exert a continuously increasing pressure on the relevant vital rate. The additional scenario of density dependence acting via recruitment limitation is closest to the case that combines (3) with (c). Obviously, the ten scenarios we consider do not represent an exhaustive list of all (infinitely many) possibilities, but are helpful for highlighting what is common and what is different between fecundity and survival regulation.

**Table 2:** Description of each of the ten scenarios used to create Figure 5.

| Scenario | Description |
| --- | --- |
| **1A** | *Exponential growth & marked excess mortality when carrying capacity is exceeded.* Standard procedure (see main text) except in generations where the pre-breeding census yields $\sum_i N_0(i) + \sum_i N_1(i) > K$, where $N_0(i)$ and $N_1(i)$ are the number of slow-senescing and fast-senescing individuals in age class $i$, and $K$ is a predefined carrying capacity. These high-density census events lead to the pre-breeding population experiencing 90% mortality across all ages of both life-history strategies (slow and fast). Thereafter, breeding and survival proceed normally (standard procedure) among the survivors. This type of regulation features exponential growth with a 'resetting' to small population sizes at regular intervals, with the cull impacting all ages equally. |
| **1B** | *Excess mortality events become more common at high density.* Like (1A), but resetting the population to small sizes occurs in a stochastic manner. Whether the 90% pre-breeding mortality occurs is decided probabilistically at each census. The probability $p$ that 90% pre-breeding mortality is applied increases with density and happens with certainty if density $K$ is reached or exceeded:<br>$p = \begin{cases} \frac{\sum_i N_0(i) + \sum_i N_1(i)}{K} & \text{if } \sum_i N_0(i) + \sum_i N_1(i) < K \\ 1 & \text{if } \sum_i N_0(i) + \sum_i N_1(i) \geq K \end{cases}$ |
| **1C** | *Continuous decline of survival with density.* At each census, we compute the value of a density-dependent factor<br>$\alpha = \begin{cases} 1 - \frac{\sum_i N_0(i) + \sum_i N_1(i)}{K} & \text{if } \sum_i N_0(i) + \sum_i N_1(i) < K \\ 0 & \text{if } \sum_i N_0(i) + \sum_i N_1(i) \geq K \end{cases}.$<br>The population follows the standard procedure as defined in the main text, but modified such that every survival value is multiplied by $\alpha$. Note that this |
|  | multiplication is also applied to juveniles born in the census year, who did not yet contribute to the census. |
| **2A** | *High density removes all old individuals.* The standard procedure is applied except when the pre-breeding census reveals a total population size above $K$, i.e. $\sum_i N_0(i) + \sum_i N_1(i) > K$. If the threshold is exceeded, all individuals above a certain age $j$, irrespective of being type 0 or 1, die before breeding begins. The value of $j$ is chosen as the largest possible age threshold (leading to the smallest possible number of age classes removed) that yields $\sum_{i=1}^j N_0(i) + \sum_{i=1}^j N_1(i) < K$, i.e. brings the pre-breeding population back to $< K$: If some individuals from age class $j$ need to be removed, everyone in that age class is removed. Afterwards, the remaining population reproduce and survive as normal (standard procedure). |
| **2B** | *High density leads stochastically to an event of removing all old individuals.* The removal event described in (2A) occurs probabilistically, with a probability that behaves like $p$ in (1B). |
| **2C** | *Continuous decline of adult survival with density.* Like (1C), but only survival of parents is negatively impacted by high density. The production and survival of newborns of the current year is unaffected, as their parents' survival is only negatively impacted after breeding occurred (Figure 4). |
| **3A** | *Crowding stops reproduction entirely.* If the census yields $\sum_i N_0(i) + \sum_i N_1(i) > K$, there is no reproduction in the given year. Extant individuals survive according to the standard procedure. |
| **3B** | *High density leads stochastically to an event of reproductive failure.* As in 3A but now a no-reproduction year occurs with probability $p$, defined as in 1A. |
| **3C** | *Continuous decline of fecundity with density.* At each census, we compute $\alpha$ as in 1C, but it is now applied to fecundities. Multiplication with $\alpha$, when performed both for $F_0$ and $F_1$, keeps the ratio $F_0:F_1$ intact. |
| **3D** | *Territoriality.* The standard procedure is applied in all other respects than juvenile survival. Adults die, which implies that there are $K - (\sum_i N_0(i) s_0(i) + \sum_i N_1(i) s_1(i))$ vacancies available once the population is proceeding towards the new census; survived juveniles are recruited to vacant territories, and should there not be sufficiently many vacancies available, the rest die. |

**Table 3:** Classification of the ten scenarios according to the type of density-dependence and according to which vital rates are affected by the density-dependence.

| Density-dependence acts on | Pulsed DD |  | Continuous DD | Competition for territories |
| --- | --- | --- | --- | --- |
|  | Deterministic | Stochastic | Deterministic |  |
| **Survival in an age-independent way** | 1A: *Exponential growth & excess mortality when carrying capacity is exceeded.* | 1B: *Excess mortality events more likely at high density.* | 1C: *Continuous decline of survival with density.* |  |
| **Adult survival in an age-dependent way** | 2A: *High density removes all old individuals.* | 2B: *High density leads stochastically to removal old individuals.* | 2C: *Continuous decline of adult survival with density.* |  |
| **Recruitment, fecundity, or newborn survival** | 3A: *Reproduction stops entirely at high density.* | 3B: *High density leads stochastically to reproductive failure.* | 3C: *Continuous decline of fecundity with density.* | 3D: *Territoriality: recruitment only possible to empty site.* |

We implement the same type of trade-off between fecundity and survival in all ten scenarios: we contrast the success of a fast life history that senesces, with a slow life history that does not experience senescence. In the above, trade-off-free section, the slow type always had an advantage, but now we switch to a trade-off: absence of senescence can only be achieved by ‘accepting’ a lower fecundity than that achieved by the fast-senescer. The ‘fast’ type thus has superior fecundity but also experiences senescence according to the Gompertz-Makeham model, where mortality has an intrinsic component that increases exponentially with age (Gompertz 1825, Makeham 1860, Missov and Lenart 2013). For simplicity, we only consider survival senescence, and we assume that fecundities do not depend on the age of the reproducing individual (while they depend on population density in three out of the ten scenarios, namely 3A-3C). We use subscript 0 to denote slow (using the mnemonic that ‘no senescence’ is indicated with a 0), and 1 denotes the fast type.

### Simulation steps shared among all scenarios

We describe here what we call the ‘standard procedure’ (Figure 4), which are the steps that are shared among all our regulation scenarios; Table 2 then describes what differs between each scenario.

Each step begins with a census of all individuals, whose ages are integers 1, 2, … with an upper limit of 200, chosen to be significantly higher than the life expectancy in any of the scenarios such that, in practice, we never observed a significant number of individuals reaching this age (we verified that the results were not changed by choosing a higher upper limit). The life cycle continues with reproduction, where slow and fast individuals’ fecundities relate to each other as a per capita fecundity ratio *F*_0_ : *F*_1_. In the standard scenario, this is achieved simply by letting slow types produce *F*_0_ offspring, while fast types produce *F*_1_. In case of fecundity regulation (three of the ten cases), the fecundities need to respond to density; we then interpret *F*_0_ and *F*_1_ as maximal fecundities in the absence of competitors, and use realized fecundities when letting strategies compete: realized fecundities are α*F*_0_ and α*F*_1_ where α < 1 takes smaller values with increasing density (Table 2).

Next in the life cycle, survival is applied deterministically, such that a proportion of individuals remain to be part of the next census (Figure 4). Survival for slow life histories equals

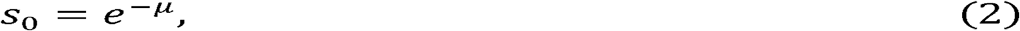

where the subscript 0 refers to slow, *μ* is extrinsic mortality (interpreted as a constant hazard, which means that 1 year is survived with *e^−μt^* where *t* = 1, hence *e^−μ^*), and there is no age-dependency since slow individuals do not senesce. Note that the ‘no age-dependency’ statement applies to the standard procedure; density-dependent adjustments may mean that survival is adjusted (multiplication with a factor α) for some age classes *s*_0_(*i*) but not others (Table 2).

Fast individuals’ survival is age-dependent to begin with (even in the standard procedure; age-dependency may become additionally modified by density dependence). In the standard procedure, we model senescence of fast individuals using the commonly used Gompertz-Makeham model of mortality which assumes mortality has a constant age-independent component μ and a component that increases exponentially with age *i*,

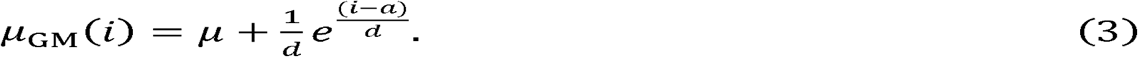

It follows that the probability that a newborn reaches age 1 (and becomes part of its first census) is 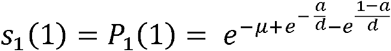. Here *s*_1_(1) denotes survival over 1 unit of time from 0 to 1, which here is the same as *P*_1_(1), the proportion of individuals still alive at age 1 (the 1 in brackets denotes age, the subscripted 1 indicates this applies to the fast strategy). For the case of newborns these are the same value (*s*_1_ (1) = *P*_1_(1)). For later ages, they are not. Generally

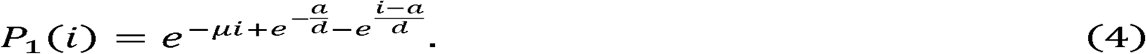

Since in our notation *s*_1_(*i*) captures survival from *i* − 1 to *i*, it equals *P*_1_(*i*)/*P*_1_(*i* − 1), which yields

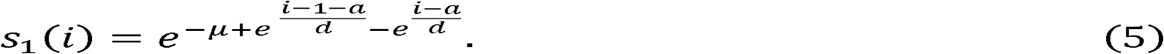

In the absence of extrinsic mortality (*μ* = 0), senescence is the only cause of death, and under this (unlikely) scenario the parameter *a* gives the modal age of death. In the presence of extrinsic mortality, *a* alone no longer translates into the modal age of death; across all values of μ ≥ 0, *a* is better interpreted as the age at which senescence acts strongly to limit lifespan (Figure 3) — loosely put, it measures how long an individual is ‘built for’. The parameter *d* impacts the variance in lifespan: at low *d* values most individuals die around the same age, at higher *d* values there is more variation in the age at death (Figure 3). As before (with *s*_0_), the values *s*_1_(*i*) can be further modified by density dependence (Table 2).

**Figure 3.**
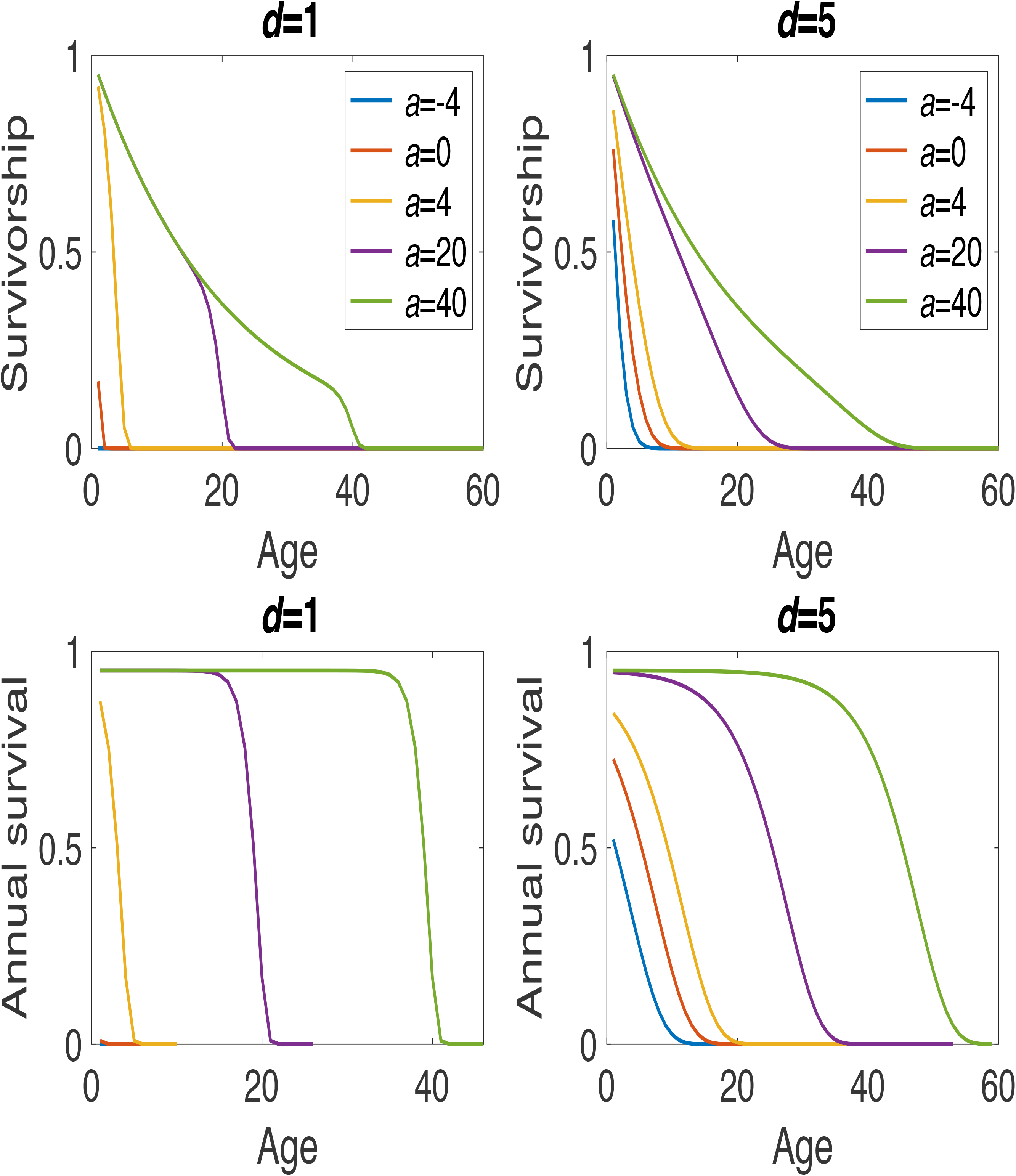
(a,b) Gompertz-Makeham survivorship function (the probability that an individual is still alive) and (c, d) age-dependent survival for example combinations of *a* and *d* as indicated. All examples use *μ* = 0.05. In the absence of extrinsic mortality (*μ* = 0), the parameter *a* gives the modal age of death. In the presence of extrinsic mortality, *a* can be interpreted as the age at which senescence starts acting strongly to limit lifespan (survivorship starts to drop rapidly around age a). The parameter *d* impacts the variance in lifespan: at low *d* values most individuals die around the same age, at higher *d* values there is more variation in the age at death.

**Figure 4:**
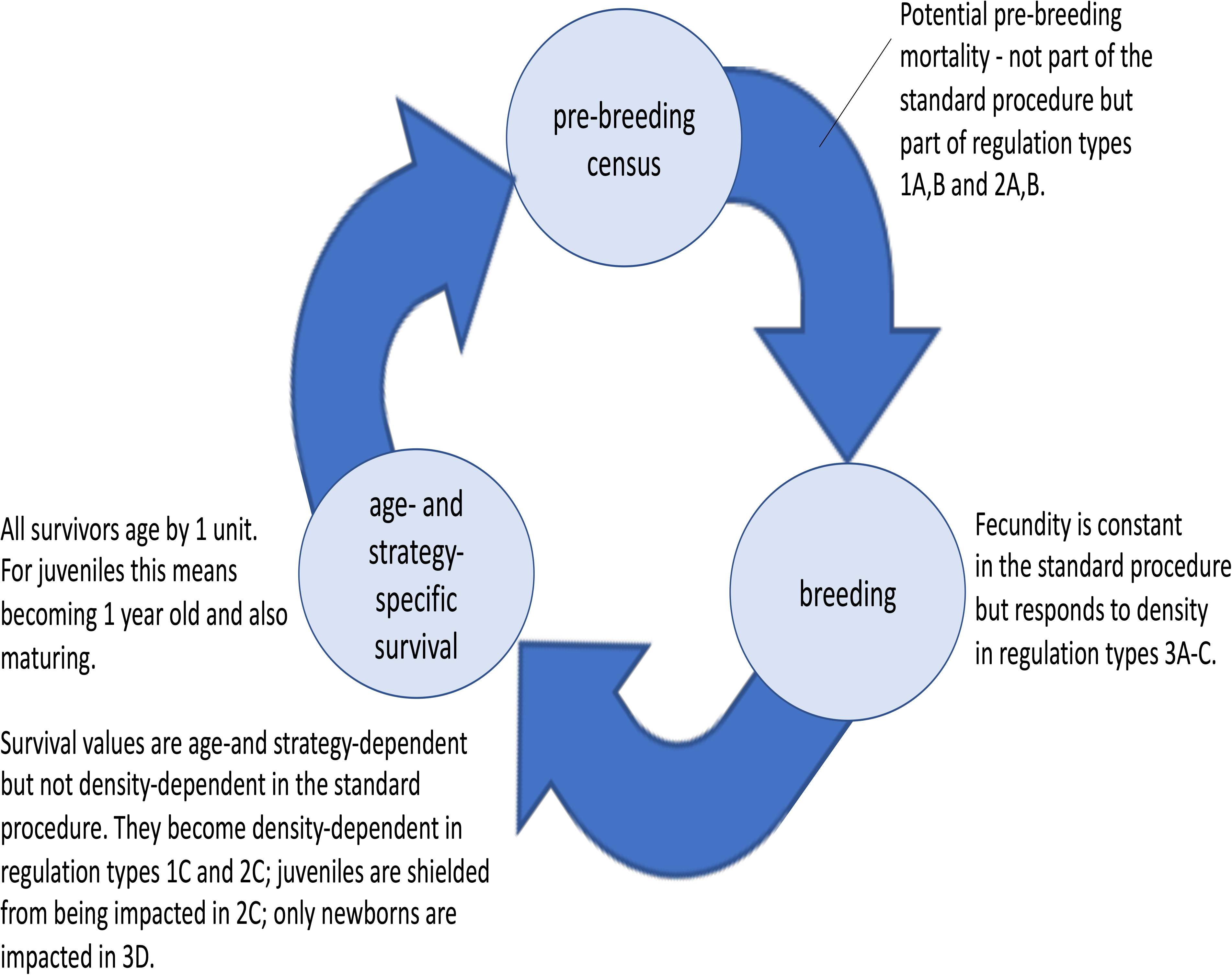
The standard procedure, describing a life cycle used to create the 10 different modes of density regulation. Note that there is no implication that completing each of the three arrows takes the same amount of time: in reality, census is immediately before breeding, to allow mortality rates time to apply before the next year. Generations are overlapping, therefore at each point the population will consist of individuals of different integer ages. Details about regulation are briefly summarized next to the loop; for full see details in Table 2.

### Model results

Clearly, we do not claim that nature offers only two life history options available for a population to choose from, or that a completely non-senescing phenotype is within the range of evolvable possibilities for many organisms (but see Roper et al. 2021 for a recent discussion on the topic). We focus on the simple contrast between an ageing high-fecundity and a non-ageing low-fecundity strategy because it serves our general aim of improving intuition about why density dependence has its known effects on the general applicability of the null result. For each of the ten types of population regulation (density dependence), we report the outcome of competition between the slow and the fast type for a range of values of extrinsic mortality *μ* (which acts on both types equally).

Whatever the value of *μ*, the outcome obviously depends on just how much lower the fecundity of the slow type is (the ratio *F*_0_:*F*_1_). Intuition suggests that there is always some intermediate value where the fates of the two strategies switch. At the one extreme, if *F*_0_ = 0, the lack of senescence of slow individuals cannot help them in competition against *F*_1_ individuals, as the former are infertile; while at the other extreme, where *F*_0_ = *F*_1_, slow individuals have a longer lifespan with no cost in fecundity, and the slow life history is guaranteed to take over. In between, there is a value of *F*_0_:*F*_1_ where selection switches from favouring fast to favouring slow. Therefore we show all results in Figure 5 in the form of an answer to the following question: what is the lowest fecundity (*F*_0_) that allows the slow strategy to outcompete the fast strategy (with fecundity *F*_1_)? And, how does this threshold depend on extrinsic mortality? If it increases with *μ*, then the ‘Williams’ prediction holds: low-*μ* conditions make it easy for slow life histories to evolve, even if building a robust body (high *a*) means sacrificing fecundity by a lot.

**Figure 5.**
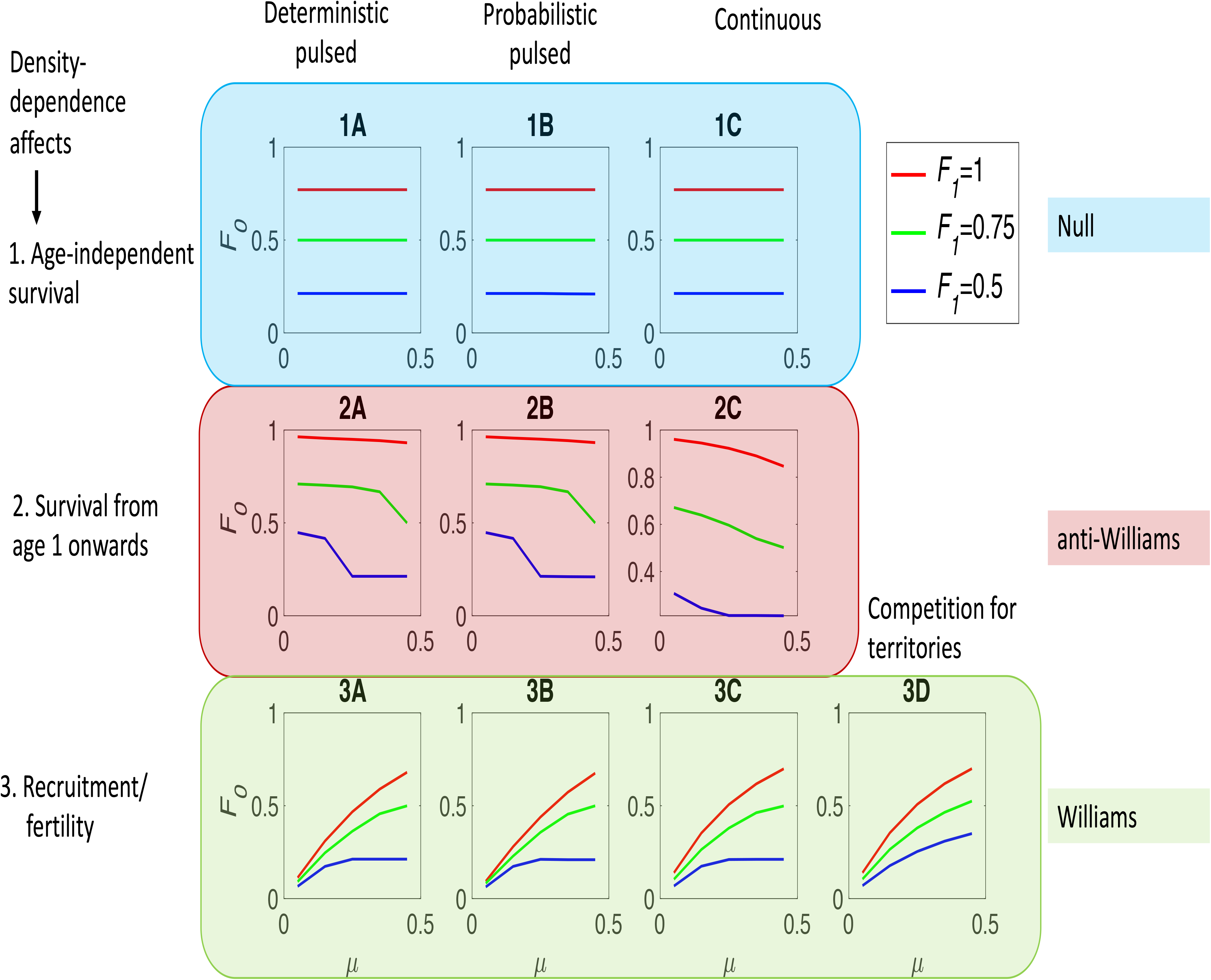
How much fertility, *F*_0_, does a non-senescing (slow) strategy need to beat the senescing (fast) strategy? The lines indicate a threshold fecundity *F*_0_ for the slow life history for three different values of fecundity of the fast strategy, *F*_1_, as indicated by the legend: above this threshold slow types win, below this threshold the fast strategy wins. Parameters used for the Gompertz-Makeham (equation 3): *a*=4, *d*=1, *K*=100 000.

The results of the ten different regulation styles are clearly categorizable in three rows (Figure 5). When density dependence causes all individuals to suffer diminished survival, extrinsic mortality has no effect on the threshold value (the threshold value of fecundity needed by the slow strategy, F_0_, to win is not affected by extrinsic mortality and therefore the line is flat in Figure 5:1A-C), i.e. the ‘null’ result holds — and it does so regardless of whether we chose a ‘pulsed’ type of regulation acting occasionally (cases 1A,B) or one of a more continuous nature (case 1C). When juveniles are shielded from the negative effects of density, however (ecologically, such a result might arise if their niche differs from that of the adults, and the adult niche is the limiting one), then an increase in extrinsic mortality makes it easier for the slow strategy to invade (the threshold value of F_0_ needed for slow to win reduces as extrinsic mortality increases leading to decreasing curves in Figure 5: 2A-2C), and we find an anti-Williams pattern. Finally, when density dependence acts on fecundity or on juvenile recruitment, an increase in extrinsic mortality makes it harder for the slow strategy to invade (the threshold value of F_0_ needed for slow to win increases as extrinsic mortality increases leading to increasing curves in Figure 5: 3A-D), in line with Williams’ hypothesis and predictions made by later models (Abrams 1993).

## Discussion

Williams’ hypothesis has triggered lively debates among theoreticians for decades. While our results do not contradict earlier work (e.g. Hamilton 1966, Charlesworth 1993, Wensink et al. 2017, Dańko et al. 2018, Day & Abrams 2020), we hope that our examples make it easier to grasp why age-independent extrinsic mortality does not affect the evolution of senescence in the absence of density-regulation, or in the presence of density-regulation that depresses survival to an equal degree across all individuals. Simultaneously, our results are fully in line with earlier findings (Abrams 1991, Day & Abrams 2020) that emphasize that Williams’ prediction is likely to hold whenever density dependence ‘hurts juveniles’ either by making it difficult for adults to produce them in the first place, or making their survival or recruitment low. There is rather broad empirical support for Williams-type patterns across species (e.g. Ricklefs 2008), which may be seen as indirect evidence that population regulation often operates via this mode.

To understand why density regulation affecting fecundity leads to a Williams-like result, it is crucial to understand that the benefits of having a robust body that can delay senescence can only materialize if the organism also avoids deaths that have nothing to do with senescence. Low extrinsic mortality means this problem is small, but do these deaths become more common as populations grow? They might not: populations can be regulated via lower juvenile production at high densities, sparing the adults. This allows the slow-senescing type to keep reaping the benefits of its robust body. The intuition that increased extrinsic mortality rate reduces the benefit of a long life is therefore correct when regulation acts on fecundity.

Our models also include the counterintuitive finding of anti-Williams patterns, where selection to avoid senescence is strongest when extrinsic mortality is high. The pattern is possible when density-dependent deaths concentrate at old ages while extrinsic mortality hits individuals regardless of their age. Under such conditions, old individuals experience both kinds of mortality risk while young ones only suffer from the extrinsic component. Thus, mortality increases with age in this setting, whatever the extrinsic mortality rate. At first sight, this appears to make investments in survival at old age unprofitable. However, the anti-Williams pattern is not defined by absolute investments, but by how the investments change with extrinsic mortality. The *relative* importance of high mortality at old age depends on extrinsic mortality: high extrinsic mortality rates imply that more deaths are age-independent while fewer are attributable to old age. At high extrinsic mortality, old individuals keep their fitness prospects reasonably intact *relative* to the prospects experienced by their younger competitors. This translates into larger benefits of having traits that combat senescence.

Our ten examples are based on a specific comparison where a no-senescing type competes with a Gompertz-Makeham-type senescer. They obviously do not constitute proof that deviant patterns could never be found, should one consider other comparisons. It is noteworthy, however, that our general statements about Williams hypothesis being strongly impacted by the mode of population regulation also apply when switching from a Gompertz-Makeham framework to survival curves that use the multi-stage model of cancer (where several stochastic ‘hits’ need to happen before the organism dies, Kokko & Hochberg 2015; code and figures at https://doi.org/10.5281/zenodo.6705180). Since different population regulation modes definitely exist (e.g., Drury & Dwyer 2005, Sæther et al. 2016, Dánko et al. 2017, Lee et al. 2021), and even a limited exploration of their consequences reveals all three patterns to exist (null, Williams and anti-Williams), it appears robust to conclude that the diversity of senescence (and lifespan) patterns across taxa are unlikely to be fully attributable to differences in extrinsic mortality alone.

Because our results show that comparative predictions ideally require an understanding of causes of shorter lifespans in one population compared with another, as well as general information on the mode of population regulation, it may be premature to make statements about individual case studies. It is nevertheless interesting that e.g. predation has been shown to impact senescence either positively or negatively. Insular populations of opossums are under lower predation pressure and senesce at a lower rate compared to mainland populations (Austad 1993). Reznick et al. (2004) on the other hand showed that guppy populations subject to *higher* predation rates senesced at lower rates than populations under lower predation risk. The latter authors speculate about possible mechanisms explaining this anti-Williams type pattern, but data is still lacking to show possible density-dependent effects on older age classes. More generally, empirical studies of the effect of extrinsic mortality on senescence usually lack evaluations of density-dependent effects on vital rates (but see Stearns et al. 2000), hindering interpretations about causal factors behind observed patterns. There is also indirect evidence for the effect of predation on senescence; patterns of senescence are compared among species with different modes of life, under the assumption that the ability to fly or to live underground decreases exposure to predation (Austad and Fischer 1991, Holmes and Austad 1994, Healy et al. 2014).

Models that include processes not included by us may highlight other reasons for finding specific patterns. Anti-Williams patterns may, for example, be found when explicitly considering condition-dependence impacting susceptibility to extrinsic mortality (the definition of ‘extrinsic’ is then subtly different: it ceases to be ‘unavoidable’ as an organism’s traits now influence its susceptibility to it). In brief, when being frail or senescent increases an organism’s susceptibility to extrinsic mortality, high extrinsic mortality leads to stronger selection on slow senescence (Abrams 1993, Williams & Day 2003). Fitting this pattern, salmon populations senesce at lower rates when predation rates by bears are high and directed towards senescing individuals specifically (Carlsson et al. 2007). Similarly, selection for heat resistance is associated with increase in lifespan in *Caenorhabditis elegans*, such that populations experiencing higher temperature-related mortality risks also senesce at slower rates (Chen and Maklakov 2012).

In some cases, there is an apparent mismatch between our predictions and data. In experimental (Stearns et al. 2000) and observational (e.g., desiccating ponds) data, high mortality or high risks of habitat disappearance are often stated to lead to faster life histories (*Daphnia*: Dudycha and Tessier 1999, killifish: Tozzini et al. 2013). Similarly, grasshoppers living at higher altitudes are subject to higher risks of freezing episodes and accordingly show faster life-histories and earlier senescence compared to populations at lower altitudes (Tatar et al. 1997). All these are examples of a Williams pattern. That this pattern arises when desiccation or freezing typically kills adults and spares eggs, may appear to be at odds with our predictions: is this not a case where we showed anti-Williams to be a possibility? The solution is to note that we did not actually model ephemeral habitats, but situations where old individuals are the first to feel effects of high density. Under ephemeral conditions, habitats are prone to disappear but this is not *caused* by high density of organisms (although *Daphnia* populations are more dense just before a desiccation event than when the hatching first began, and there may be more grasshoppers late in the season than early, this is correlation, not causation: an abundance of *Daphnia* does not cause ponds to dry and winter does not happen because grasshoppers became abundant). Had we modelled ephemeral habitats but not (yet) incorporated any causal link from high density to poor performance, we would not have had a mechanism in place that prevents unbounded growth. For the population to be regulated, some additional causality will need to be incorporated, and a modeller will still have the freedom to choose which part of the population will suffer. Exactly how these may restore a Williams pattern remains an open question; models of this type should probably also consider that ephemeral habitats may cut individual lives short before maturity is reached. Speeding up maturation time may be an adaptive response in such situations, with effects felt throughout the life cycle.

There is a strong parallel between the theoretical discussions surrounding the Williams hypothesis and the debate that has arisen in the study of within-species patterns of humans specifically (Frankenhuis & Nettle 2020, Dinh et al. 2022, Sear 2020, Stearns & Rodrigues 2020, Woodley of Menie et al. 2021). Whether environmental harshness selects for faster life-history strategies is much debated in this field (del Giudice 2020, Frankenhuis & Nettle 2020, Galipaud & Kokko 2020, Sheehy-Skeffington 2020, Lynch et al. 2020, Dinh et al. 2022, Zietsch & Sidari 2020). Our results suggest that understanding population regulation is crucial in this context too (see also Baldini 2015). Human populations have clearly experienced rather different growth conditions over their existence (examples: Hamilton et al. 2007, 2009, de Pablo et al. 2019, Bird et al. 2020, Freeman et al. 2020), and it is not *a priori* clear what is a typical enough pattern to have been the relevant selective environment. To what extent within-species plasticity can be assumed to be aligned with evolutionary predictions derived for whole populations is likewise debated (del Giudice 2020, Galipaud & Kokko 2020 and references therein). Note that in our models we assumed the latter, i.e. evolutionary responses of entire populations, as there were no environmental cues that individuals could perceive and change their location on the slow-fast continuum accordingly. Actual models of plastic responses are rare for human life histories (Nettle et al. 2013, Frankenhuis et al. 2018), and it would be interesting to see if their predictions change as dramatically with population regulation modes as our current results show to be the case with evolutionary responses.

Finally, we have intentionally chosen to model trade-offs in a stylized way, leaving subtleties such as the difficulty of optimizing function simultaneously for early and late life (Maklakov & Chapman 2019) for later studies. Our flat, decreasing and increasing curves in Figure 5 correspond to null, anti-Williams and Williams, respectively, in a simple setting where a slow pace of reproduction makes organisms immune to senescence. We fully admit that our assumptions are difficult to interpret in cases where high densities remove the oldest age classes: how can an organism combine a lack of senescence with higher vulnerability to high density at old age? Future work is clearly needed, with more explicit links between performance at early and later ages, perhaps with a mechanistic focus, explicit models for gene expression, or e.g. the system reliability approach (Gavrilov & Gavrilova 2001, Laird & Sherratt 2009, 2010a,b) as well as selection that relates to the possibility of indeterminate growth (Vaupel et al. 2004, Caswell & Salguero-Gómez 2013, Purchase et al. 2022). While the multitude of factors listed above suggest that wide diversity in senescence patterns and lifespans (Jones et al. 2014) is the expectation, we hope that our conceptual examples help to see why a specific feature of life cycles – the diversity in modes of population regulation — continue to play a very important role.

## Data, scripts and codes availability

Matlab scripts are online: https://doi.org/10.5281/zenodo.6705180.

## Acknowledgements

We thank the PCI recommender Sinead English, reviewer Nicole Walasek and one anonymous reviewer for comments that helped improve our manuscript. We also thank Willem Frankenhuis and Daniel Nettle for encouragement and discussions.

